# Autophagic degradation of the Cucumber mosaic virus virulence factor 2b balances antiviral RNA silencing with proviral plant fitness and virus seed transmission

**DOI:** 10.1101/2020.02.13.938316

**Authors:** Aayushi Shukla, Gesa Hoffmann, Silvia López-González, Daniel Hofius, Anders Hafrén

## Abstract

Autophagy is a conserved intracellular degradation pathway that has recently emerged as an integral part of plant responses to virus infection. The elucidated mechanisms of autophagy range from the selective degradation of viral components to a more general attenuation of disease symptoms. In addition, several viruses are able to manipulate the autophagy machinery and counteract autophagy-dependent resistance. Despite these findings, the complex interplay of autophagy activities, viral pathogenicity factors, and host defence pathways in disease development remains poorly understood. In the current study, we analysed the interaction between autophagy and Cucumber mosaic virus (CMV) in *Arabidopsis thaliana*. We show that autophagy is induced during CMV infection and promotes the turnover of the major CMV virulence protein and RNA silencing suppressor 2b. Intriguingly, 2b itself dampens plant autophagy. In accordance with 2b degradation, we found that autophagy provides resistance against CMV by reducing viral RNA accumulation in an RNA silencing-dependent manner. Moreover, autophagy and RNA silencing pathways contribute to plant longevity and fecundity of CMV infected plants in an additive manner, uncoupling it from resistance. In addition to reduced fecundity, autophagy-deficient plants also failed to support seed transmission of the virus. We propose that autophagy attenuates CMV virulence via 2b degradation and thereby increases both plant and virus fitness with a trade-off penalty arising from increased RNA silencing-mediated resistance.

**Author summary:** The capacity of plants to fight pathogenic viruses in order to survive and minimize damage relies on profound cellular reprogramming events. These include the synthesis of new as well as the degradation of pre-existing cellular components, together shifting cellular homeostasis towards a better tolerance of disease and fortification of antiviral defence mechanisms. Autophagy is a prominent and highly conserved cellular degradation pathway that supports plant stress resilience. Autophagy functions vary broadly and range from rather unspecific renewal of cytoplasm to highly selective degradation of a wide collection of specific substrates. Autophagy is well established to be involved during virus infections in animals, and its importance has also recently emerged in virus diseases of plants. However, we are still far from a comprehensive understanding of the complexity of autophagy activities in host-virus interactions and how autophagy pathway engineering could be applied against viruses. Here, we have analyzed one of the traditional model plant viruses, Cucumber mosaic virus (CMV), and its interactions with autophagy. Our study revealed that autophagy is tightly integrated into CMV disease, influencing processes from plant health to CMV epidemiology.

## Introduction

Autophagy is a conserved eukaryotic mechanism that is important for cellular homeostasis through its functions as a major catabolic system. In its simplest form, the autophagy process is considered to sequester cytoplasmic portions non-selectively within a vesicular structure called the autophagosome, which subsequently enters the lytic vacuole of plants for degradation and recycling of nutrients. However, the extent to which autophagy operates in a non-selective manner still remains an open question. In parallel, the examples of selective targeting of autophagic substrates by specialized cargo receptors continues to increase (1, 2). The plant autophagic molecular machinery was initially characterized in *Arabidopsis thaliana* and subsequently recognised in algae, gymnosperms and other angiosperms (3). Core components of this pathway include numerous ATG genes, such as ATG5, ATG7 and ATG8. Selective autophagy is known to take part in the turn-over of substrates including chloroplasts, peroxisomes, aggregates, ribosomes and proteasomes in plants. Selectivity opens the possibility for several highly distinct autophagic processes to operate in parallel, and outlines the important challenge to dissect the overall autophagy effect into its finer mechanisms.

Autophagy is induced by numerous environmental conditions, including abiotic stress and nutrient limitation. Its importance for plant adaptation is established in part by the increased sensitivity of autophagy-deficient plants to these conditions (3, 4). Autophagy also plays important roles in plant-pathogen interactions including diseases caused by viruses, bacteria, fungi and oomycetes (5). These studies support the notion that autophagy is a complex process with several functions operating in parallel to influence the infection outcome. Because pathogens commonly cause severe stress to their host plants, it is not surprising that infected plants show an upregulation of autophagy upon infection (6–10). This finding outlines the important concept that the interaction between pathogens and their hosts likely co-evolves in the presence of an activated autophagy response. As a consequence, both pathogen and host could try to utilize the plant autophagy process for their own benefit.

In virus-plant interactions, we can distinguish between two mechanisms that limit disease. Host resistance refers to the ability to suppress virus multiplication whereas tolerance is defined as the ability of the plant to minimise the infection-associated damage as a result of the pathogen infection. Co-adaptation of hosts and viruses has resulted in a range of interactions between antiviral resistance mechanisms and viral counter-strategies, together balancing plant and virus fitness (11). Autophagy has been established as a prominent response pathway to several viral infections in the animal field (12). An emerging theme from these studies is that viruses have acquired properties to modify autophagy in many ways, including induction, suppression and subversion of its functions. Evolution of such viral properties can be considered to directly reflect the importance of autophagy for virus infection. Recently, fundamental roles for autophagy have been identified in plant viral diseases, including autophagic degradation of viral components, virus-based counteraction and modulation of autophagy, as well as autophagy-mediated promotion of plant tolerance (7–9, 13, 14). Interestingly, a connection between autophagy and the foremost antiviral defence pathway in plants, RNA silencing, is emerging. Initially, autophagy was suggested to degrade the RNA silencing suppressors of potyviruses (HCpro) and cucumoviruses (2b) (15), and we could recently show that the autophagy cargo receptor NBR1 degrades HCpro to reduce virus susceptibility and accumulation (7). Additionally, autophagy degrades the satellite ßC1 RNA silencing suppressor of geminiviruses, AGO1 when targeted by the poleroviral RNA silencing suppressor P0, and SGS3/RDR6 in the presence of potyviral VPg (9, 16, 17). Notably, autophagy-based resistance was uncoupled from autophagy-based tolerance against a *Potyvirus* (Turnip mosaic virus; TuMV) and a *Caulimovirus* (Cauliflower mosaic virus; CaMV) (7, 8). Thus, autophagy shows complex interactions and a general potential for regulating plant virus disease.

In this study we show that autophagy is induced during infection by Cucumber mosaic virus (CMV; genus *Cucumovirus*), a positive-stranded RNA virus. We found that the viral RNA silencing protein 2b has a modest capacity to dampen autophagy and is itself subject to autophagic degradation. Because autophagy suppressed CMV RNA accumulation in an RNA silencing- and AGO1-dependent manner, we propose that 2b degradation represents a resistance mechanism that sensitizes CMV to RNA silencing. In a broader context, autophagy and RNA silencing appear beneficial for CMV infection through synergistic promotion of plant longevity, fecundity and viral seed transmission. Our results thereby reveal that autophagy provides resistance by reducing virus accumulation as well as tolerance by decreasing disease severity during CMV infection, and both processes seem to be linked to the autophagic degradation of the major virulence factor 2b.

## Results

### Autophagy suppresses disease severity and CMV accumulation

A potential aspect influencing virus and plant fitness is the severity of disease. Previously, we showed that biomass loss, as a measure for disease, was severely increased in loss-of-function mutants of the core autophagy gene ATG5 but not in plants lacking the autophagy cargo receptor NBR1 when infected with TuMV or CaMV (7, 8). In this study, we analysed whether CMV-infected plants showed a similar dependence on functional autophagy by infecting WT, *atg5* and *nbr1* plants with CMV and scoring disease symptoms at 28 days after infection (DAI). While infected WT and *nbr1* plants appeared indistinguishable (Fig 1A) and showed a similar loss in fresh weight (Fig 1B), infected *atg5* plants were severely dwarfed and showed a higher fresh weight loss. We conclude that the core autophagy machinery, but not NBR1 is important for the plant to attenuate CMV disease. This observation is similar to previous findings for TuMV and CaMV (7, 8), and supports a universal role for autophagy in reducing disease symptoms of pathogenic plant viruses.

**Fig 1.**
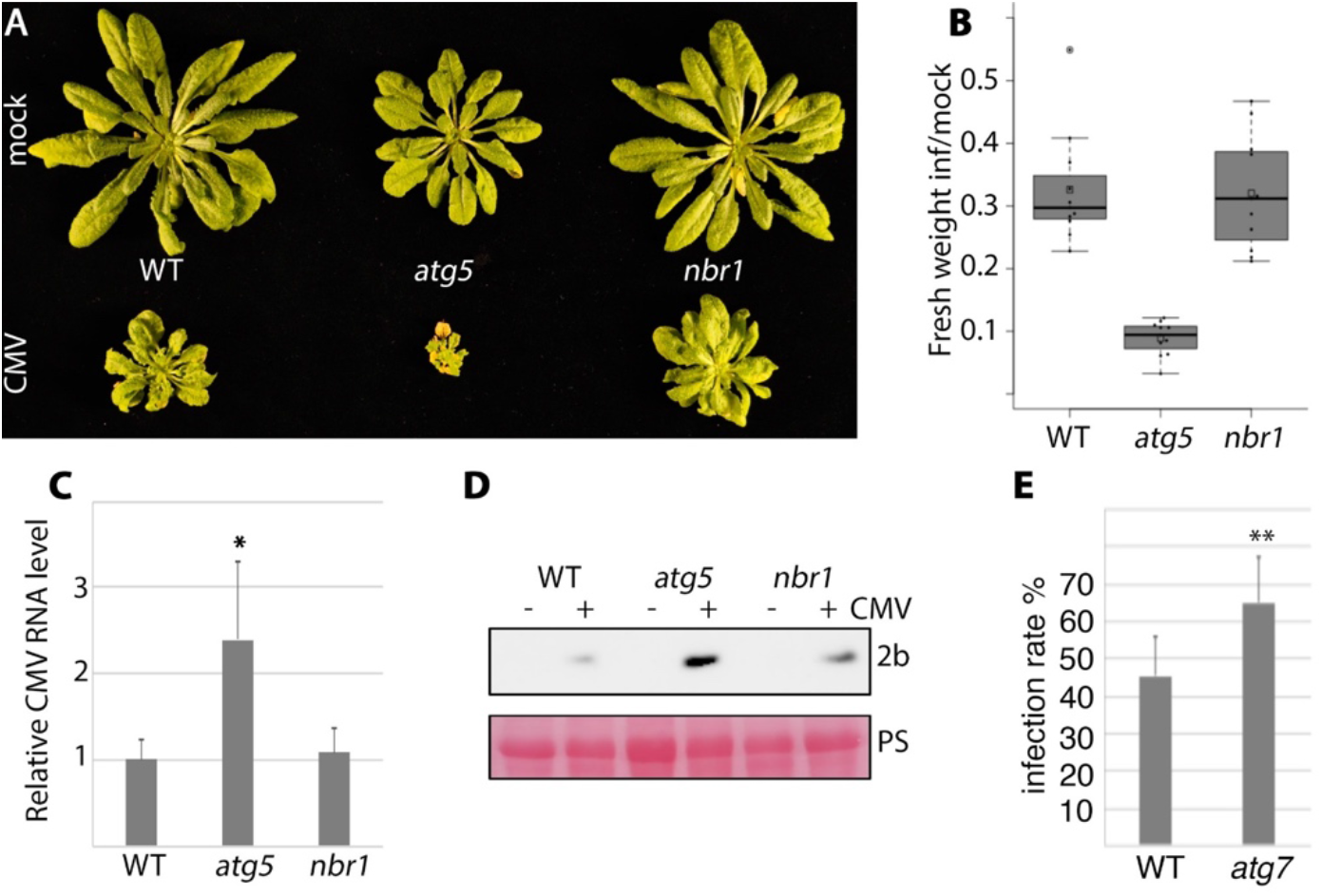
Autophagy promotes plant tolerance and resistance against CMV infection. A) Representative WT (Col-0), *atg5* and *nbr1* plants at 28 days after mock or CMV inoculation. B) The fresh weight ratio of CMV infected to mock plants 28 days after inoculation (DAI). (*n*=9). C) Relative CMV RNA levels determined by RT-qPCR at 14 DAI. (*n*=4). D) In parallel with (C), protein samples were prepared and CMV protein 2b was detected by Western blotting using anti-2b. NBR1 detection was used to verify *atg5* and *nbr1* mutants and Rubisco large subunit was visualized by Ponceau S staining (PS) of the membrane as loading control. E) CMV infection rate in WT and *atg7* plants determined by presence/absence of viral symptoms three weeks after mechanical sap inoculation. (*n*=8). Statistical significance (\**P*<0.05; \*\**P*<0.01) was revealed by Student’s *t*-test (compared to WT).

Next, we determined CMV multiplication in WT, *atg5* and *nbr1* plants at 14 DAI (Fig 1C). Viral RNA accumulated to higher levels in *atg5* compared to WT and *nbr1* plants. Notably, autophagy-dependent suppression of CMV RNA accumulation was NBR1-independent, distinguishing it from selective autophagy pathways contributing to resistance against CaMV and TuMV (7, 8). We could also detect higher levels of the viral 2b protein in *atg5* compared to WT and *nbr1* plants by Western blotting (Fig 1D), further supporting that autophagy limits CMV accumulation independently of NBR1. We further estimated virus susceptibility by calculating the number of plants showing infection symptoms after mechanical sap inoculation of CMV, and found that infection rate was increased in autophagy-deficient *atg7* mutants compared to WT plants (Fig 1E). Together, these results establish autophagy as a suppressor of CMV infection by decreasing susceptibility, viral RNA accumulation and disease symptomology.

### Autophagy is induced during CMV infection

Owing to the effect of autophagy in CMV infection (Fig 1), we set out to determine whether autophagy levels were altered during infection. We used an Arabidopsis line that stably expresses GFP-ATG8a, which is a commonly applied marker for detection of autophagosomes. Distribution of this marker was altered in CMV infected tissue, including increased numbers of smaller GFP-ATG8a puncta that likely represented autophagosomes (Fig 2A and B). Infected tissue also contained larger GFP-ATG8a labelled structures that were not observed in healthy tissue. Treatment with concanamycin A (conA), an inhibitor of the vacuolar ATPase, was used to stabilize autophagic bodies delivered into the vacuole (8). The number of GFP-ATGA8a puncta substantially increased upon conA application in infected tissue (Fig 2A and B), suggesting that autophagy is both activated and completed during CMV infection.

**Fig 2.**
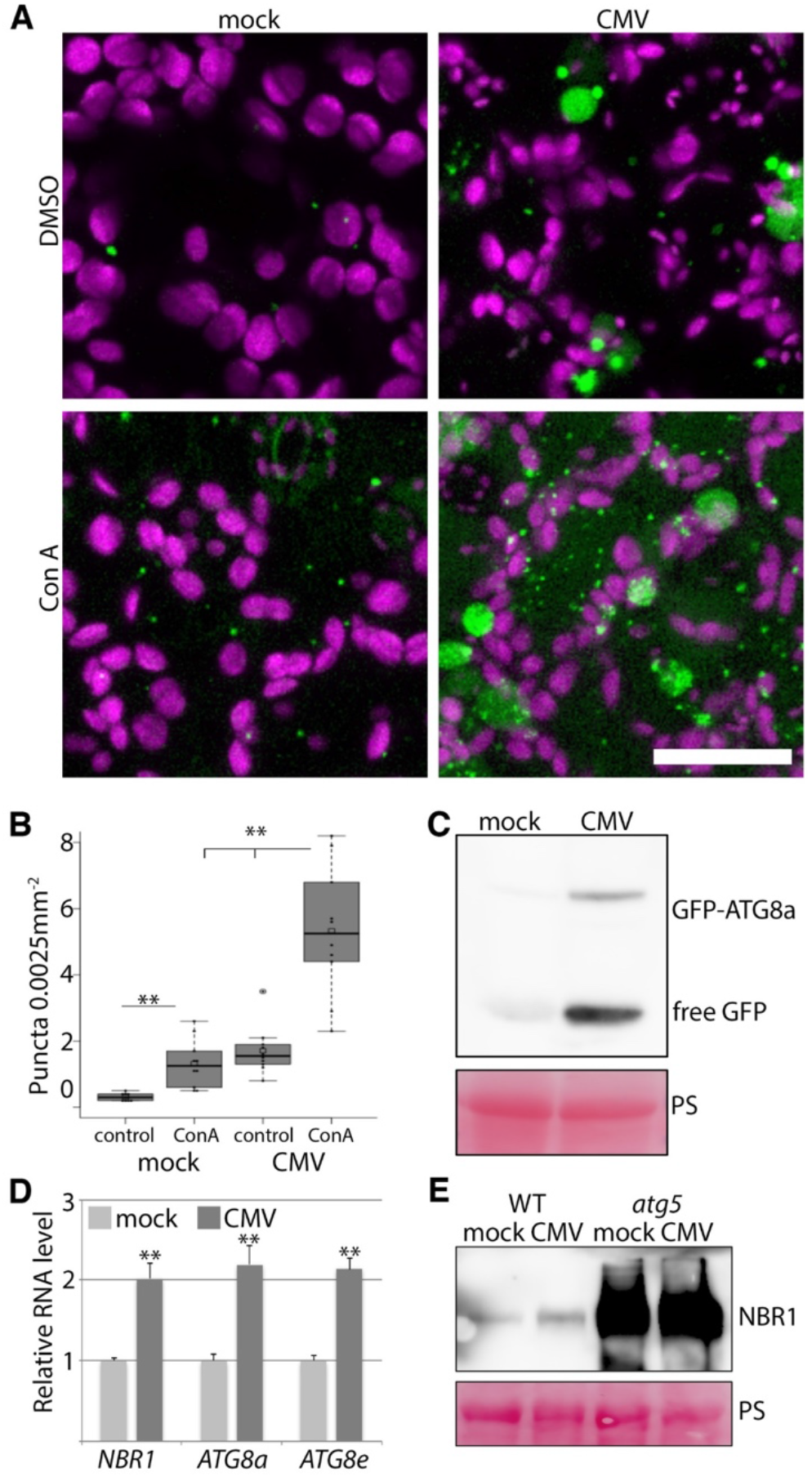
Autophagy is activated and functional during CMV infection. A) Representative images of the GFP-ATG8a marker in healthy and CMV infected plants with and without concanamycin A (conA) treatment. Chloroplasts are shown in magenta. Images are confocal Z-stacks. Scale bar = 20 μm. B) GFP-ATG8a foci were counted from similar images as in (A) using Image J. (*n*=10). C) Western blot analysis of free GFP levels derived from GFP-ATG8a in mock and CMV infected plants. Ponceau S (PS) staining verified comparable protein loading. D) Transcript levels for *NBR1*, *ATG8a* and *ATG8e* in mock and CMV infected plants were determined by RT-qPCR. (*n*=4). E) Western blot detection of NBR1 levels in mock and CMV inoculated WT and *atg5* plants. Ponceau S (PS) staining verified comparable protein loading. Statistical significance (\**P*<0.05; \*\**P*<0.01) was revealed by Student’s *t*-test (compared to WT).

The processing of free GFP from the GFP-ATG8 fusion protein is regarded as another proxy for autophagy flux in plants. Free GFP levels were higher in CMV infected tissue compared to the healthy control (Fig 2C), supporting elevated autophagy levels. Moreover, we observed higher transcript levels of autophagy-related genes *ATG8a*, *ATG8e* and *NBR1* in CMV infected tissue (Fig 2D), suggesting transcriptional activation of autophagy. NBR1 delivers cargo to autophagosomes and gets degraded in the process, making the NBR1 protein to mRNA ratio another indicator of autophagy activity and flux. Notably, despite elevated *NBR1* transcript levels, there was only slight increase in NBR1 protein accumulation (Fig 2E), suggesting enhanced autophagic degradation during CMV infection. These assays together indicated that autophagy is induced and functional during CMV infection.

### CMV 2b protein modulates the autophagy response

Our observation of infection-induced autophagy together with the autophagy-dependent suppression of viral RNA accumulation and disease severity prompted us to analyse whether individual CMV proteins could modulate autophagy. First, we used a recently developed quantitative assay in *Nicotiana benthamiana* leaves, that is based on the transient expression of ATG8a fused to *Renilla* luciferase (RLUC) together with firefly luciferase (FLUC) as internal control (6). Co-expression of the viral protein 2b substantially increased RLUC-ATG8a accumulation in relation to FLUC, while 2a, 3a and CP behaved similar as the GUS control (Fig 3A). This result identified 2b as a potential suppressor of autophagy, and as none of the viral proteins seemed to recapitulate infection-induced autophagy on their own, this elevation is likely to be a result of the full infection process. Over-expression of ATG3 was previously shown to induce autophagy in *N. benthamiana* (18) and indeed resulted in a 5-fold increase of FLUC to RLUC-ATG8a ratio (Fig 3B). Additional expression of 2b diminished the inducing effect by ATG3, further supporting the autophagy-suppressing activity of 2b in this system.

**Fig 3.**
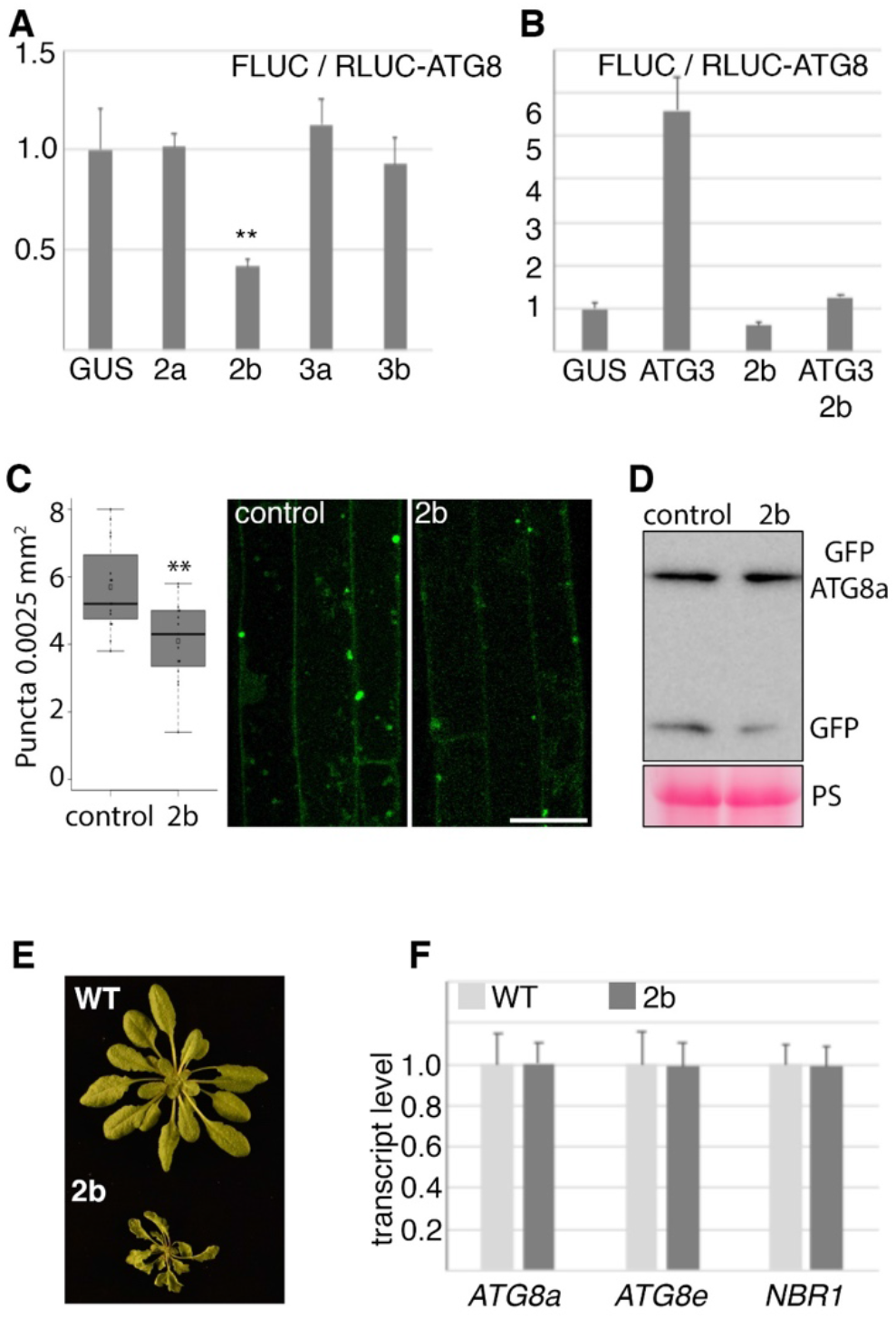
CMV protein 2b can dampen autophagy. A) RLUC-ATG8a and the internal control FLUC were co-expressed with viral proteins and GUS in leaves of *N. benthamiana*. Normalized luciferase values are given as a ratio between FLUC and RLUC, resulting in decreased ratios when RLUC-ATG8a degradation is reduced. (*n*=4). B) As in (A), but co-expressed proteins where GUS, ATG3, 2b and ATG3+2b. (*n*=4). C) The number of GFP-ATG8a foci were analyzed in roots of 10-day old seedlings without and with stable expression of 2b after 10h conA treatment. Counting was performed using ImageJ particle analyzer on confocal Z-stack projections. (*n*=15). Representative images are shown with scale bar = 20 μm. D) Anti-GFP western blot analysis was performed in parallel with (C) to estimate the ratio of GFP-ATG8a to free GFP. Rubisco large subunit visualized by Ponceau S staining (PS) of the membrane is shown as loading control. E) Representative image of 4-week-old short-day grown 2b expression line in comparison to WT plants. H) Transcript levels of *ATG8a*, *ATG8e* and *NBR1* were determined by RT-qPCR using *PP2a* as reference from plants shown in (E). (*n*=4). Statistical significance (\*\**P*<0.01) was revealed by Student’s *t*-test.

Importantly, we also found that autophagy was modestly down-regulated in 2b transgenic Arabidopsis seedlings as indicated by a reduced number of GFP-ATG8a labelled autophagosomes in conA treated seedling roots (Fig 3C) and a decreased accumulation of free GFP from GFP-ATG8a (Fig 3D). 2b is the major pathogenicity factor of CMV and 2b transgenic plants show severe growth retardation as well as different morphological abnormalities (19) (Fig 3E). However, the transcript levels for *ATG8a*, *ATG8e* and *NBR1* remained unaffected in 2b transgenic plants (Fig 3F). Thus, the 2b-mediated dampening of autophagy levels appear to occur post-transcriptionally and, 2b pathogenicity alone is not causative for the up-regulation of autophagy pathway transcripts during CMV infection (Fig 2D). We therefore conclude that 2b is capable of attenuating the autophagy response.

### CMV 2b is degraded through autophagy

Previously, autophagy-dependent 2b degradation has been proposed based on its accumulation in response to the PI3K inhibitor 3-methyladenin during transient 2b expression in *N. benthamiana* (15). The marked accumulation of the CMV RNA silencing suppressor 2b in *atg5* (Fig 1E) supported this notion that 2b can be degraded by autophagy. Additionally, we also found 2b accumulation in the *atg7* mutant compared to WT (Fig 4A). To further support 2b by autophagy, we performed a GFP-based immunoprecipitation of GFP-ATG8a from mock and CMV infected plants. We found that 2b, but also viral CP, co-purified with GFP-ATG8a (Fig 4B), suggesting both proteins may be degraded by autophagy during infection.

**Fig 4.**
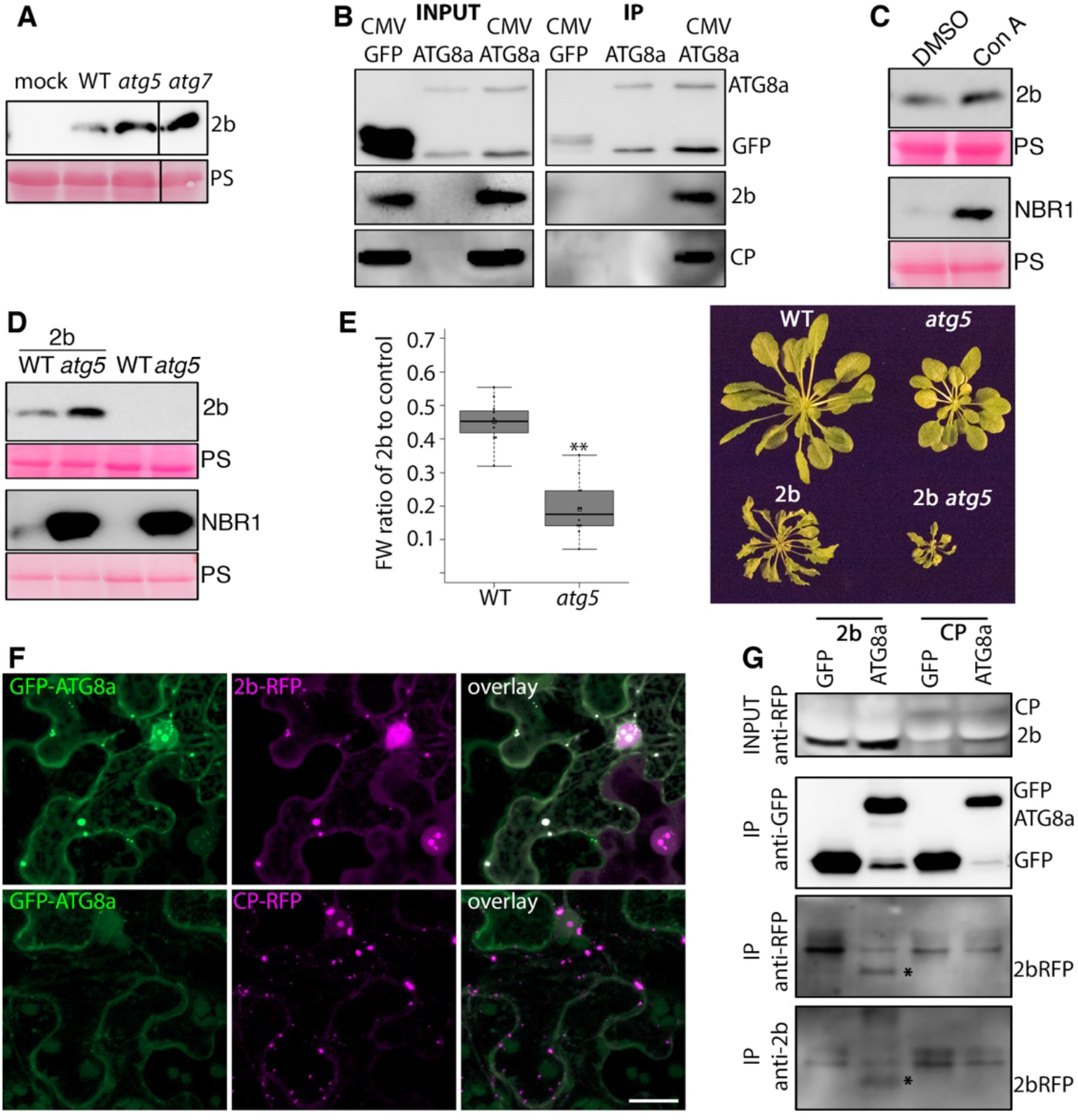
Autophagy degrades CMV 2b both in and outside the infection context. A) Accumulation of viral 2b protein was determined in WT, *atg5* and *atg7* plants at 14 DAI by western blot analysis. B) Co-immunoprecipitation analysis of viral CP and 2b with GFP-ATG8a from infected transgenic Arabidopsis tissue. Non-infected GFP-ATG8a and infected GFP expressing plants were used as control. Shown are the input and immunoprecipitated (IP) samples. C) 2b transgenic seedlings were treated with DMSO (control) or concanamycin A for 10h followed by western blot detection of 2b. Increased accumulation of NBR1 verified the concanamycin A (conA) treatment. D) Accumulation of 2b in transgenic WT and *atg5* plants was analyzed by western blotting. Increased accumulation of NBR1 verified the *atg5* background. Ponceau S-stained Rubisco shows loading (PS) in (A, C and D). E) 2b-dependent virulence measured as relative fresh weight loss caused by transgenic 2b expression compared to non-transgenic plants in WT and *atg5* backgrounds. (*n*=9). A representative plant image is shown to the right. F) Co-localization analysis in *N. benthamiana* leaves co-expressing 2b-RFP or CP-RFP with GFP-ATG8a. These Z-stack images were acquired 48 h post agroinfiltration. Scale bar = 20 μm. G) Co-precipitation analysis of 2b-RFP and CP-RFP with GFP-ATG8a in parallel with (F). Expression of GFP was used as control. Shown is the anti-RFP input signal, as well as anti-GFP, anti-RFP and anti-2b signals from the GFP-based immunoprecipitated samples. The position of 2b-RFP is marked with an asterisk.

We further sought to analyse the autophagy-dependent targeting of 2b outside the infection context. We found only a minor stabilization of 2b by conA in transgenic 2b seedlings (Figure 4C), while NBR1 accumulated to substantial amounts, thus verifying the successful inhibitor treatment. We speculated that either the rate of 2b autophagic degradation is slower than that of NBR1 or the stability of these proteins differ in the vacuole. Nonetheless, after introgression of the 2b transgene into the *atg5* background, we found much stronger accumulation of 2b in *atg5* compared to WT (Fig 4D). Notably, NBR1 accumulated to higher levels in WT plants expressing 2b, further supporting 2b-mediated reduction of autophagy. We also noticed that the relative decrease in rosette biomass caused by 2b was much higher in *atg5* plants compared to WT (Fig 4E), which could be a consequence of higher 2b levels, as 2b was shown to facilitate virulence in a concentration-dependent manner (19), and increased sensitivity of *atg5* to 2b virulence.

Finally, we carried out co-localization and immunoprecipitation experiments addressing 2b and CP association with ATG8a. 2b-RFP, but not CP-RFP, co-localized with GFP-ATG8a in larger cytoplasmic structures (Fig 4F). 2b-RFP was also associated with multiple nuclear speckles including the previously described targeting to the nucleolus (20), and caused re-localization of GFP-ATG8a to the same domains (S1 Fig). CP-RFP also localized to the nucleolus but without recruiting GFP-ATG8a. Interestingly, 2b re-localized CP-GFP but not free GFP to the nuclear sub-compartments during their co-expression (S1 Fig). When 2b-RFP and CP-RFP were co-expressed with either GFP or GFP-ATG8a followed by GFP-based immunoprecipitation, we could detect specific association of 2b with ATG8a. An unspecific signal arising from the RFP-antibody obscured judgement of co-purification of viral CP. Also, no signal was detected in the purified samples using an antibody against CP (not shown), and together with the absence of co-localisation between CP and ATG8a (Figure 4F), CP appears not to be a prominent autophagy target, at least outside of infection. In conclusion, 2b protein associates with ATG8a inside and outside of an infection context.

### Suppression of CMV RNA accumulation by autophagy involves AGO1 and Dicer proteins

CMV 2b suppresses DCL2- and DCL4-dependent antiviral RNA silencing as shown by restored pathogenicity of 2b-deficient CMV in the *dcl2 dcl4* knock-out mutant (21, 22). Furthermore, 2b interacts directly with AGO1 as part of RNA silencing suppression (23). We hypothesized that autophagy could suppress CMV RNA accumulation by compromising 2b-dependent suppression of RNA silencing. If this was the case, autophagy-dependent suppression of CMV RNA accumulation should be less pronounced in *dcl2 dcl4* and *ago1* backgrounds. Indeed, while CMV RNA accumulated to higher levels in both *atg7* and *dcl2 dcl4* mutants, there was no additive effect in the *atg7 dcl2 dcl4* background at 14 DAI (Fig 5A). We obtained similar results for AGO1 as the *ago1 atg5* double mutant did not show any additive increase in CMV RNA accumulation compared to *atg5* and *ago1* single mutants (Fig 5B). The absence of additive effects supports that autophagy and AGO1/DCL2,4-dependent resistance are coupled, likely by autophagy targeting of the viral RNA silencing suppressor 2b.

**Fig 5.**
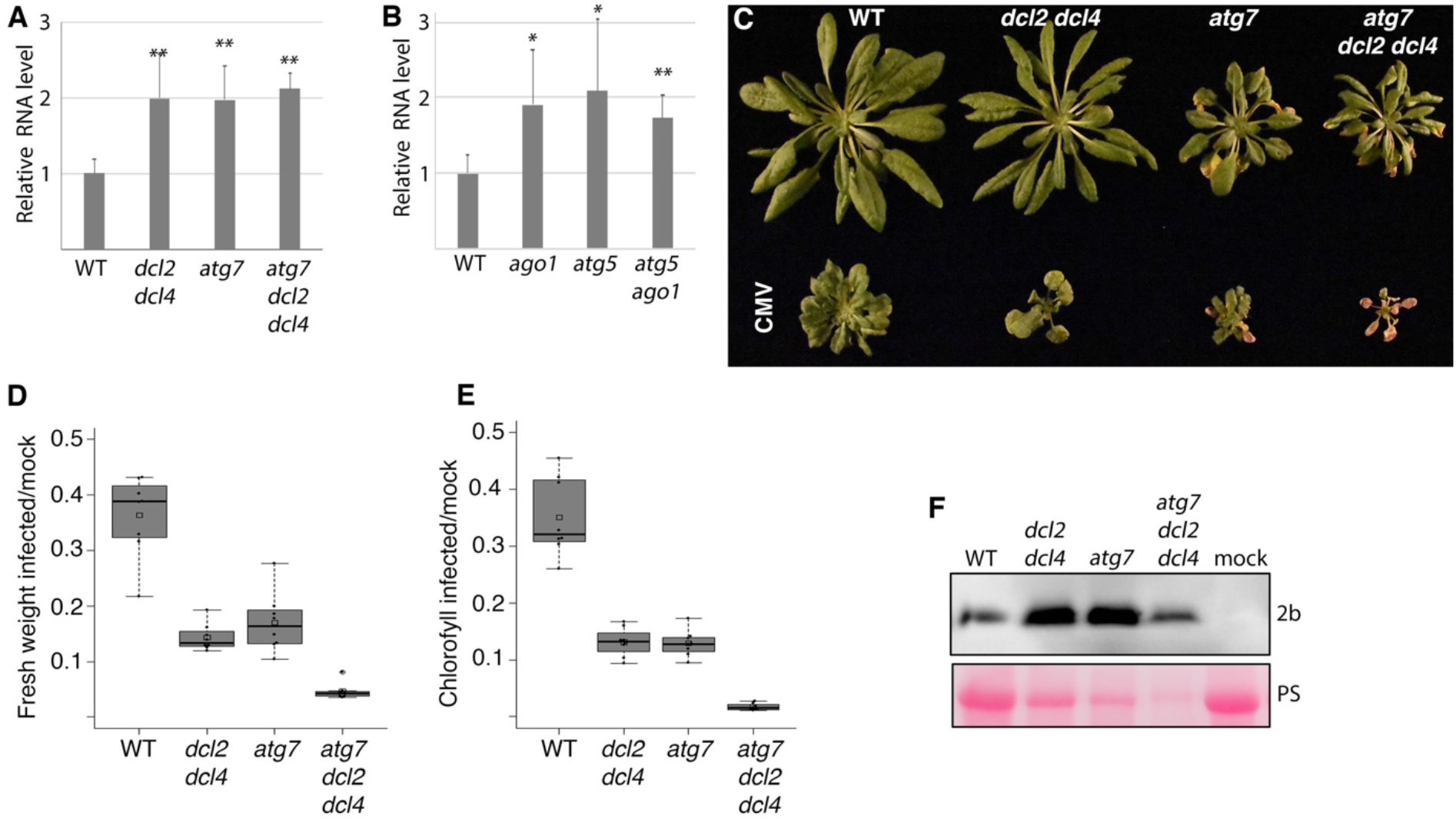
Autophagy-enhanced CMV resistance and tolerance are uncoupled and involve RNA silencing. A) Relative CMV RNA levels determined by RT-qPCR at 14 DAI in WT, *atg7*, *dcl2 dcl4* and *atg7 dcl2 dcl4* plants (*n*=4). B) Relative CMV RNA levels determined by RT-qPCR at 14 DAI in WT, *atg5*, *ago1* and *atg5 ago1* plants (*n*=4). C) Representative image of mock-and CMV inoculated plants at 28 DAI. D and E) Relative fresh weight (D) and chlorophyll content (E) in CMV infected plants compared to mock at 28 DAI. (*n*=10). F) Western blot analysis of CMV 2b protein accumulation per plant in parallel with (C, D, E). Ponceau S stained Rubisco shows loading (PS) and reveals reduced total protein content per plant in the mutants compared WT.

### Autophagy and RNA silencing synergistically support plant tolerance to CMV

Despite the apparent non-additive functions of autophagy and RNA silencing in reducing CMV RNA accumulation at early stage of infection (Figure 5A and B), *atg7 dcl2 dcl4* mutants eventually showed a total collapse when infections progressed further (Fig 5C). Notably, there was no tissue senescence in either WT or *dcl2 dcl4* plants, while this autophagy-associated phenotype was clearly visible in CMV-infected *atg7* and highly intensified in *atg7 dcl2 dcl4* plants. When the fresh weight of the rosette was compared as a proxy of disease severity between WT, *atg7*, *dcl2 dcl4* and *atg7 dcl2 dcl4* plants at 28 DAI, a strong additive effect on biomass loss was observed in the combinatorial mutant of autophagy and RNA silencing (Fig 5D). We detected a similar dependence on biomass loss between *atg7* and *dcl2 dcl4* for the unrelated dsDNA virus CaMV but not the RNA virus TuMV (S2 Fig). Possibly, TuMV fails to show this dependence owing to very strong suppression of RNA silencing, as biomass loss was overall highest for TuMV and comparable between WT and *dcl2 dcl4* in contrast to CaMV and CMV. The severe biomass loss during CMV infection in the *atg7 dcl2 dcl4* mutants was further corroborated by the loss of chlorophyll (Fig 5E). Because of the severe disease phenotype at 28 DAI, we chose to analyse the amount of viral 2b protein per plant as a measure of virus accumulation instead of RT-qPCR. 2b accumulated to clearly higher levels in the single mutants compared to WT (Fig 5F). 2b accumulated to higher levels also in the triple mutant compared to WT when related to the loading control, but whether there are additive effects between *atg7* and *dcl2 dcl4* remained unclear. These results suggested that impaired resistance could contribute to the severe disease phenotype in the *atg7* and *dcl2 dcl4* mutants, and we speculate that the escalated disease phenotype in the triple mutant may be a combination of the failure to undergo partial recovery in *dcl2 dcl4* and enhanced virulence sensitivity in *atg7*.

### Autophagy is essential for transmission of CMV through seeds

CMV belongs to those plant viruses that also transmit trans-generationally through seeds in a process termed as vertical transmission. Vertical transmission is correlated with plant fecundity, which we found to be substantially more reduced in autophagy-deficient *atg7* compared with WT plants (Fig 6A). Interestingly, CMV infected *dcl2 dcl4* plants failed to produce seeds. Despite reduced fecundity, the seed germination potential remained comparable between *atg7* and WT irrespective of CMV infection (Fig 6B). Finally, we estimated CMV seed transmission in offspring seedlings from infected WT and *atg7* plants. In WT plants, CMV showed a transmission frequency of 1.31 ± 0. 8% when calculated using Gibbs and Gower formulae (Fig 6C). This frequency of seed transmission is in the same range that has been reported for other CMV strains in Arabidopsis (24). Notably, we did not detect any seed transmission in *atg7* plants, establishing autophagy as essential for vertical transmission of CMV strain PV0187 in Arabidopsis.

**Fig 6.**
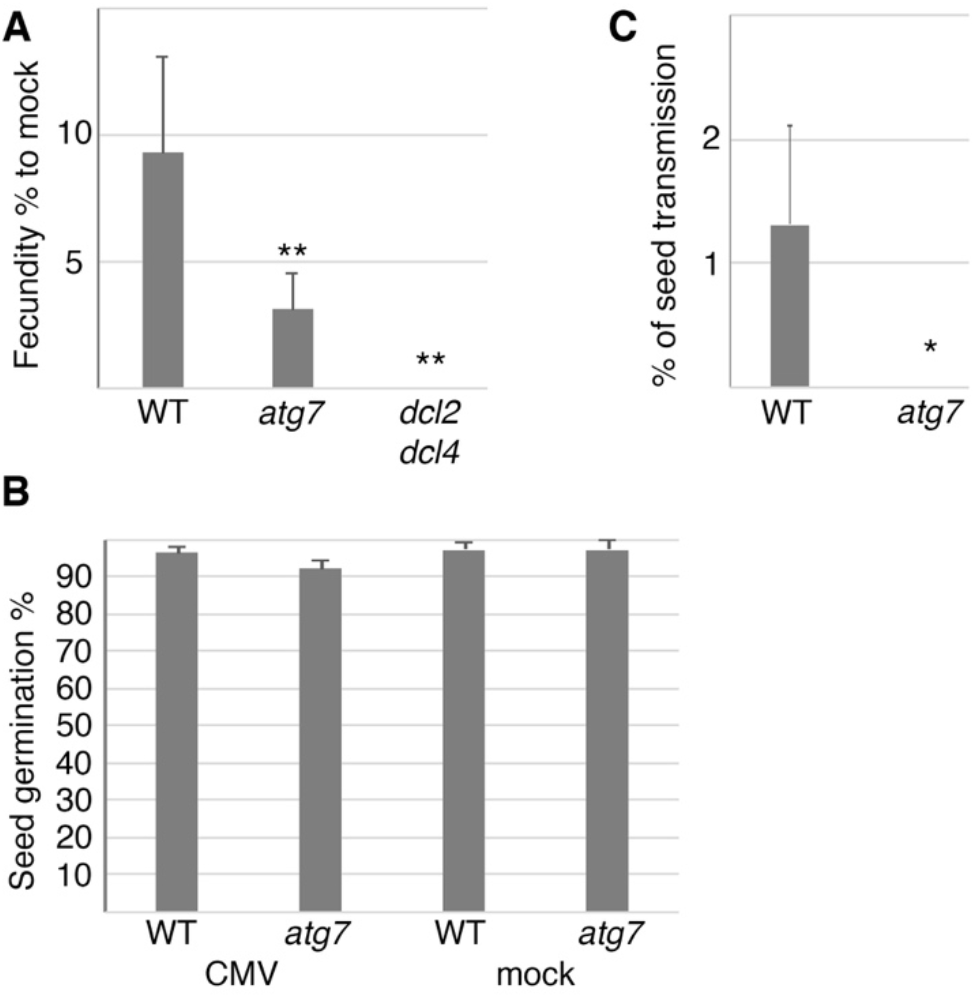
Autophagy and RNA silencing is essential for vertical transmission of CMV. A) Total seed weight of infected WT, *atg7* and *dcl2 dcl4* plants presented as a percentage relative to seeds produced by non-infected mock plants. (*n*=5). B) The percentage of seeds germinating from infected and non-infected WT and *atg7* plants. (*n*=5). C) Vertical transmission of CMV was estimated by RT-PCR in seedlings derived from seeds of infected WT and *atg7* plants (*n*=5 independent plants).

## DISCUSSION

Autophagy can deliver a wide array of substrates including proteins, RNAs, ribosomes, proteasomes, viral particles, organelles and aggregates for degradation in the lytic vacuole (1). Hence, it plays an important role in maintaining plant homeostasis, especially when plants encounter stressful conditions such as viral infections (25). The complexity of substrates targeted by autophagy underscore the possibility that several distinct autophagic processes operate in parallel upon cellular reprogramming and adaption to new conditions. This is fortified by e.g. the uncoupling of autophagy-based plant virus resistance and tolerance against TuMV (7), CaMV (8) and as demonstrated by this study, also CMV. The interaction between CMV and autophagy is complex. It includes resistance and disease attenuation that is coupled to RNA silencing components DCL2/DCL4 and AGO1 as well as autophagy-based degradation of the major virulence factor 2b. Interestingly, 2b alone reduces plant growth in an autophagy-dependent manner and shows a moderate capacity to dampen the autophagy response. The most striking phenotype is the severely increased virulence of CMV during autophagy deficiency that results in reduced plant life span, seed production and vertical virus transmission. Taken together, these multiple connections between autophagy and infection emphasise the complexity of this interaction in CMV epidemiology.

Plants reduce pathogen virulence by tolerance and resistance mechanisms, which are assumed to impose different selective pressures on both pathogens and hosts (26). In this study we found that autophagy contributes to the resistance against CMV by reducing virus accumulation. While autophagy-mediated resistance has been established for several animal viruses (12), comparative findings were largely lacking for plant viruses. Only very recently we and others have found that autophagy targets multiple plant viruses, including the positive-stranded RNA viruses TuMV and Barley stripe mosaic virus (7, 14, 27), the negative-stranded RNA virus Rice stripe virus (13), the double-stranded DNA virus CaMV (8) and three single-stranded DNA geminiviruses (9). These findings established autophagy as a central component of plant immunity against viruses, which is fortified further by our current results on CMV. Notably, these studies have revealed a wide mechanistic diversity of autophagy-based virus resistance and viral counter-strategies (25), which ultimately suggests that different plant viruses have co-evolved with the autophagy pathway in an individual manner. However, despite the profound mechanistic differences in the detailed interaction between individual viruses and autophagy, a common theme arising is the involvement of the RNA silencing pathway in the autophagy-virus interplay (25). This connection is evident for CMV as we show that autophagy degrades the viral silencing suppressor 2b and that the enhancement of viral RNA accumulation in autophagy- and RNA silencing-deficient plants lack epistasis. These findings suggest that autophagy-based CMV resistance requires a functional RNA silencing pathway, and we speculate that this is causally linked to the degradation of the viral silencing suppressor 2b. Notably, 2b is a highly complex pathogenicity factor that among other functions interacts with AGO1/4 to impair slicing and also binds siRNAs directly (20, 23, 28, 29). Previously we showed that the TuMV silencing suppressor HCpro is degraded by autophagy, also resulting in reduced virus accumulation (7). Intriguingly, this effect appears to include the degradation of potyvirus-induced structures reminiscent of RNA granules to which also AGO1 localizes (30). Together with the autophagic degradation of the geminiviral satellite ßC1 silencing suppressor (9), autophagy appears to be more generally nested into the antiviral RNA silencing defense with virus-specific adaptations. Opposite to these antiviral roles of autophagy, viral proteins have also been proposed to utilize autophagy for degrading the RNA silencing components SGS3 and AGO1 (16, 17). Taking this into account, we need to consider that autophagy could degrade 2b alone but also in complex with siRNA, AGO1 and other components. At least for HCpro, autophagy seems to target higher-order RNA-protein complexes reminiscent of virus-induced RNA stress granules and processing bodies (7, 30) with likely complex consequences on infection. Whether viral proteins have evolved to function as autophagy cargo receptors to mediate degradation of RNA silencing components is indeed an exciting option. Our results, however, do not support that 2b uses this strategy to counteract RNA silencing-dependent resistance, and in general autophagy seems to support rather than antagonize resistance against plant viruses (7–9, 13, 14, 27).

Perhaps the most striking observation in the context of CMV and other viral infections is the escalated disease severity in autophagy-deficient plants (Fig 5). It is likely that elevated 2b levels contributes to this phenotype as 2b virulence is dose-dependent (19) and accordingly the growth of 2b transgenic *atg5* is severely reduced compared to 2b transgenic control plants (Fig 4). Defects in RNA silencing alone also increase CMV virulence, but when combined with autophagy deficiency the virulence intensifies to the extent that plants collapse and thus terminate infection. Theory suggests that under strict vertical transmission, virulence should be negatively correlated with the transmission rate (31–33). This can be reasoned because viral fitness is tightly linked to the reproductive success of the host (34). Evidently, optimized virulence benefits virus epidemiology, where virus accumulation and virulence need to be balanced with plant longevity and fecundity to promote both successful horizontal and vertical virus transmission (24, 32, 35). Thus, as autophagy and RNA silencing enhance plant longevity, fecundity and vertical transmission of CMV, the antiviral nature of these pathways in a more holistic and epidemiological context becomes less clear. 2b of CMV subgroup I, including the virulent strains PV0187 (this study) and FNY, but not the mild LS strain of subgroup II, localizes to the nucleolus (20, 36). Interestingly, nucleolar 2b appears to promote virulence uncoupled from virus accumulation and this virulence is furthermore dampened by the RNA silencing pathway (36). This finding is similar to our observation in combined autophagy and RNA silencing knockout mutants. In this context, it is notable that 2b re-localized ATG8a to the nucleolus. Functions associated with nuclear and nucleolar localized LC3, the mammalian homolog of ATG8, are still undefined (37), and so far ATG8 has not been reported in the nucleolus of plants. Despite a potential connectivity between these events, whether 2b-dependent virulence involves nucleolar ATG8 functions remains a matter of future study.

2b is a major virulence factor and RNA silencing suppressor of CMV (36) and the strong penalty of super-virulence arising from autophagy- and RNA silencing-deficiency highlights the importance for CMV to fine-tuning these interactions. One exciting possibility is that CMV recruits the autophagy pathway to degrade 2b in a regulated manner to minimize the resistance penalty and support long-term tolerance. At the plant level, virus infections are prolonged processes and the absolute amount of virus can continue to increase for several weeks during systemic infection. However, the active cellular infection cycle is assumed to be largely completed within 24 hours for most plant viruses, including CMV (38). During this relatively short period, CMV may benefit from dampening the autophagy response using 2b without any long-term tolerance costs. It is also interesting that the infection frequency is elevated in autophagy-deficient plants for CMV (Fig 1), TuMV and CaMV (7, 8). This suggests that autophagy suppresses initiation of plant virus infections in general, manifesting the autophagy-virus interaction as most decisive at the very early infection stage. Once the active stage of infection ceases in a cell, the autophagy-mediated clean-up of virulence factors like 2b should promote long-term tolerance and possibly even reduce co-infection competition from weaker viruses that would benefit from high levels of 2b. In line with this, the CP of CMV was recently proposed to destabilize 2b in a self-attenuation mechanism by which the virus achieves long-term virulence reduction (39). At this point, we can only speculate if CP-mediated 2b degradation involves autophagy, but it is interesting that both 2b and CP were present in GFP-ATG8a co-immunoprecipitations from infected plants (Fig 4) and that both proteins localize to the nucleolar virulence domain of 2b (S2 Fig) (36). Nevertheless, our results revealed the autophagic degradation of 2b during transgenic expression in the absence of CP, showing that CP is at least not strictly required in this process. Taken together, we provide a hypothetical model of autophagy as a complex regulator of CMV infection that influences virus accumulation, transmission and plant disease (Fig 7).

**Fig 7.**
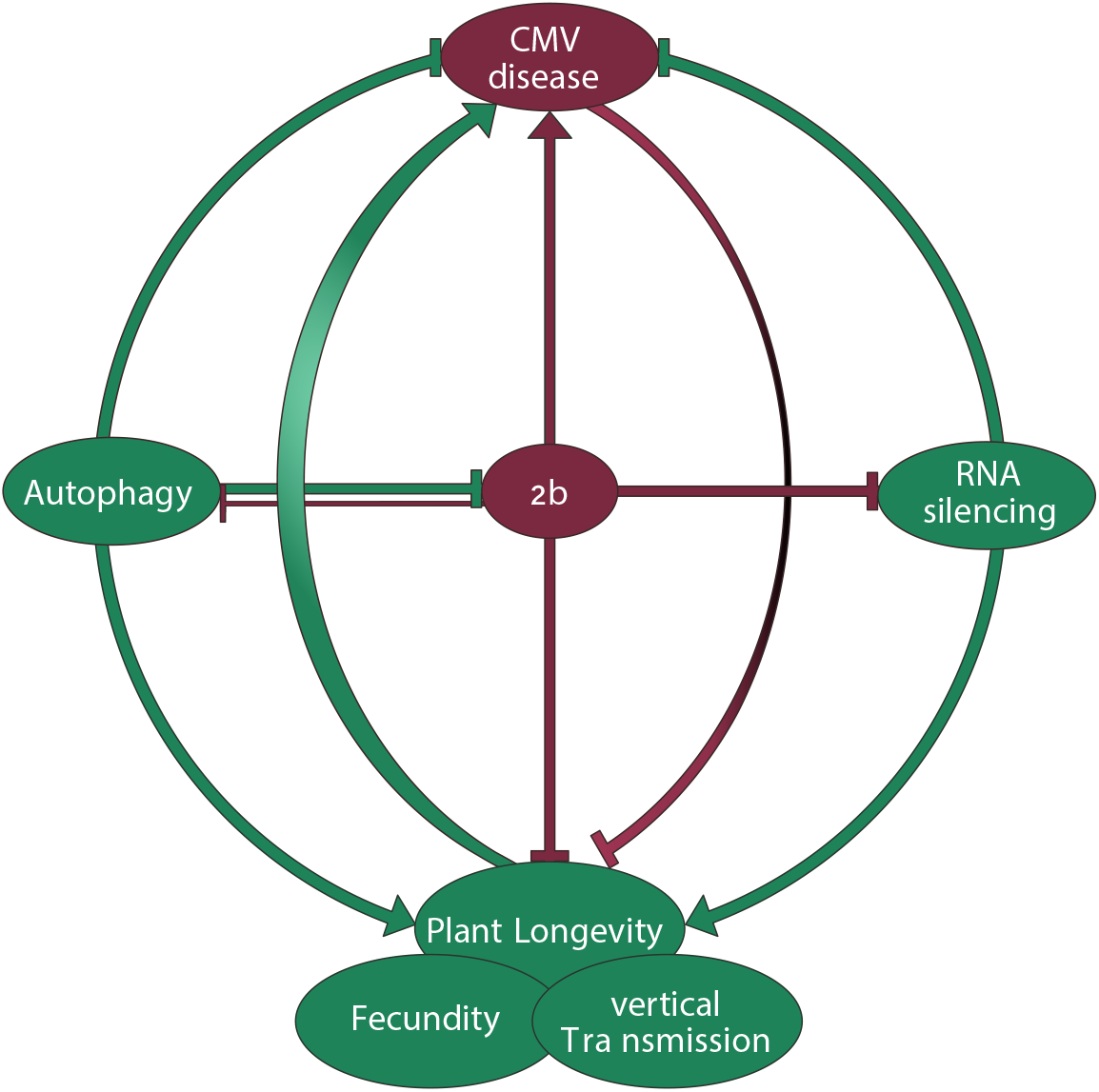
Interplay between autophagy, RNA silencing and 2b shape CMV disease. The 2b protein is the major pathogenicity determinant in CMV infection. Plant autophagy suppresses CMV accumulation through 2b degradation. By limiting 2b levels, autophagy relaxes 2b mediated suppression of antiviral RNA silencing and virulence. Vice versa, 2b itself has the capacity to restrict the plant autophagy response, but its significance during infection remains to be resolved. Both autophagy and antiviral RNA silencing suppress virus accumulation, and the lack of additivity between the pathways suggests their strong interaction in the process, potentially through 2b. Taken together with their prominent synergism in disease attenuation, we propose that autophagy and RNA silencing-based plant tolerance is not quantitatively coupled to virus accumulation and that the pathways rather operate in a parallel manner to promote survival of infected plants. Virulence evolution and trade-offs are complicated for pathogens that utilize both vertical and horizontal transmission. Autophagy and RNA silencing promotes CMV vertical transmission and likely also vector-mediated horizontal transmission through increased plant longevity with only minor fitness penalties on virus accumulation. We consider that CMV has adapted to and benefits from these potential antiviral pathways. Thus, the interplay between the viral 2b protein, plant autophagy and RNA silencing pathways determines the delicate balance between virus accumulation, transmission and plant fitness in CMV disease.

## Materials and methods

### Plant material and growth conditions

Wild-type (WT) plants were *Arabidopsis thaliana* ecotype Columbia (Col-0). Mutants *atg5-1*, *atg7-2*, *nbr1-2*, *ago1-27*, *dcl2 dcl4, atg7 dcl2 dcl4*, and the GFP-ATG8a transgenic line have been described previously (40–43). The *ago1-27 atg5-1* double mutant was generated by crossing. The transgenic line 2b3C was described previously (19), and used for crossing with GFP-ATG8a and the *atg5-1* background. Arabidopsis plants were grown on soil for infection experiments under short-day conditions (8/16-h light/dark cycles) in a growth cabinet, and *Nicotiana benthamiana* plants were cultivated for transient expression assays under long-day conditions (16/8-h light/dark cycles) in a growth room at 150 μE/m^2^s, 21°C, and 70% relative humidity, respectively.

### DNA constructs

*ATG3* and *ATG8a* as well as viral cistrons *2a, 2b, 3a* and *CP* were amplified using cDNA prepared from CMV infected plant total RNA as template and cloned into pENTRY-Topo. *ATG3* was further recombined into pGWB614, and viral proteins into pGWB660 (44). Expression constructs for GUS and GFP-ATG8a were described in (8) and pRD400::*FLUC* in (45). *ATG8a* was recombined into pMDC32:*RLUC* (6) and all binary vectors were transformed into *Agrobacterium* C58C1 for transient expression in *N. benthamiana* or transformation of Arabidopsis by the floral dip method.

### CMV inoculation and quantification

The first true leaves of 3-week-old Arabidopsis plants were inoculated mechanically with sap prepared from *N. benthamiana* plants infected with the CMV strain PV0187 described in (46). Plants were sampled in biological replicates, each containing 3 individual plants from which inoculated leaves were removed. For CMV RNA or plant transcript quantitation, total RNA was isolated using the RNeasy Plant Mini Kit (Qiagen), and on-column DNA digestion was performed with DNase I (Qiagen). First-strand cDNA was done using Maxima First Strand cDNA Synthesis Kit (Thermo Fisher Scientific). Quantitative RT-PCR analysis (qPCR) was performed with Maxima SYBR Green/Fluorescein qPCR Master Mix (Thermo Fisher Scientific) using the CFX Connect^TM^ Real-Time PCR detection system (BIO-RAD) with gene-specific primers listed in Supplemental Table 1. Normalization was done using *PP2A* (*AT1G69960*).

### Fresh weight and chlorophyll analysis

Fresh weight refers to the areal rosette of Arabidopsis plants after infection or stable transformation, and is presented as ratios to the respective mock or non-transformed plants. The relative chlorophyll content of plants was determined as described before (8).

### Confocal microscopy and inhibitor treatment

Live cell images were acquired from abaxial leaf epidermal cells using a Zeiss LSM 780 microscope. Excitation/detection parameters for GFP was 488 nm/490-552 nm. Inhibitor treatment was carried out with 0.5 μM concanamycin A in ^1^/ MS 10 h before confocal analysis. Confocal images were processed with ZEN (version 2011). Quantitation of GFP-ATG8a labelled puncta was done using Image J (version 1.48v) software essentially as described in (7).

### Immunoblot analysis

Proteins were extracted in 100 mM Tris pH 7.5 with 2% SDS, boiled for 5 min in Laemmli sample buffer and cleared by centrifugation. The protein extracts were then separated by SDS-PAGE, transferred to polyvinylidene difluoride (PVDF) membranes (Amersham, GE Healthcare), blocked with 5% skimmed milk in PBS, and incubated with primary antibodies anti-NBR1 (47), anti-2b (20), anti-CP (Bioreba Art.Nr. 160612), anti-GFP (Santa Cruz Biotechnology; sc-9996) using 1:2000 dilution in PBS 0.1% Tween-20, and secondary horseradish peroxidase-conjugated antibodies 1:10 000 in PBS 0.1% Tween-20 (Amersham, GE Healthcare). The immunoreaction was developed using the ECL Prime kit (Amersham, GE Healthcare) and detected in a LAS-3000 Luminescent Image Analyzer (Fujifilm, Fuji Photo Film).

### Immunoprecipitation

For immunoprecipitation, plant tissue was homogenized in 2 ml buffer (100 mM Tris pH8, 150 mM NaCl, 0.5% TX-100, protease inhibitor cocktail (Roche)) per gram of tissue. The lysate was cleared at 4000 x g for 5 min at 4°C, filtered through two layers of miracloth and incubated 1h with anti-GFP μbeads according to manufacturer’s instruction (Milteney) for infected Arabidopsis tissue and GFP-agarose beads (Chromotech) for transiently expressing *N. benthamiana* tissue. After washing 4 times with buffer, samples were eluted using 2x Laemmli sample buffer and analyzed by immunoblotting.

### Analysis of fecundity, germination potential and vertical virus transmission

Plants grown under long-day conditions were infected with CMV and allowed to set seeds. The seed weight was recorded and the relative fecundity is presented as a ratio between seeds from control and infected plants. Seed germination potential of offspring seeds was recorded after calculating the percentage of germinated seeds on ½ MS plates supplemented with 1% sucrose. For vertical transmission, 10 pools of 20 seedlings each were analysed by RT-PCR per parental plant for presence of CMV. Probability of virus transmission by a single seed was calculated by the Gibbs and Gower’s formulae (48) 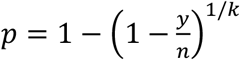; where p is the probability of virus transmission by a single seed, y is the number of positive samples, n denotes the total number of samples assayed and k represents the number of seedlings taken per sample.

### Data analysis and presentation

Data are presented as mean ± standard deviation (SD) and statistical significance was analyzed by two-sided Student’s *t*-test with *p*-values <0.05 denoted * and *p*-values <0.01 denoted **. The number of replicates is given in the respective figure legends (*n*).

## Acknowledgements

We gratefully acknowledge John Carr for providing the CMV PV0187 strain and 2b transgenic lines as well as Thomas Canto for the 2b antibody. Funding from FORMAS (grant number 2016-01044) for A.H. is also gratefully acknowledged.

---

**S1 Fig.**
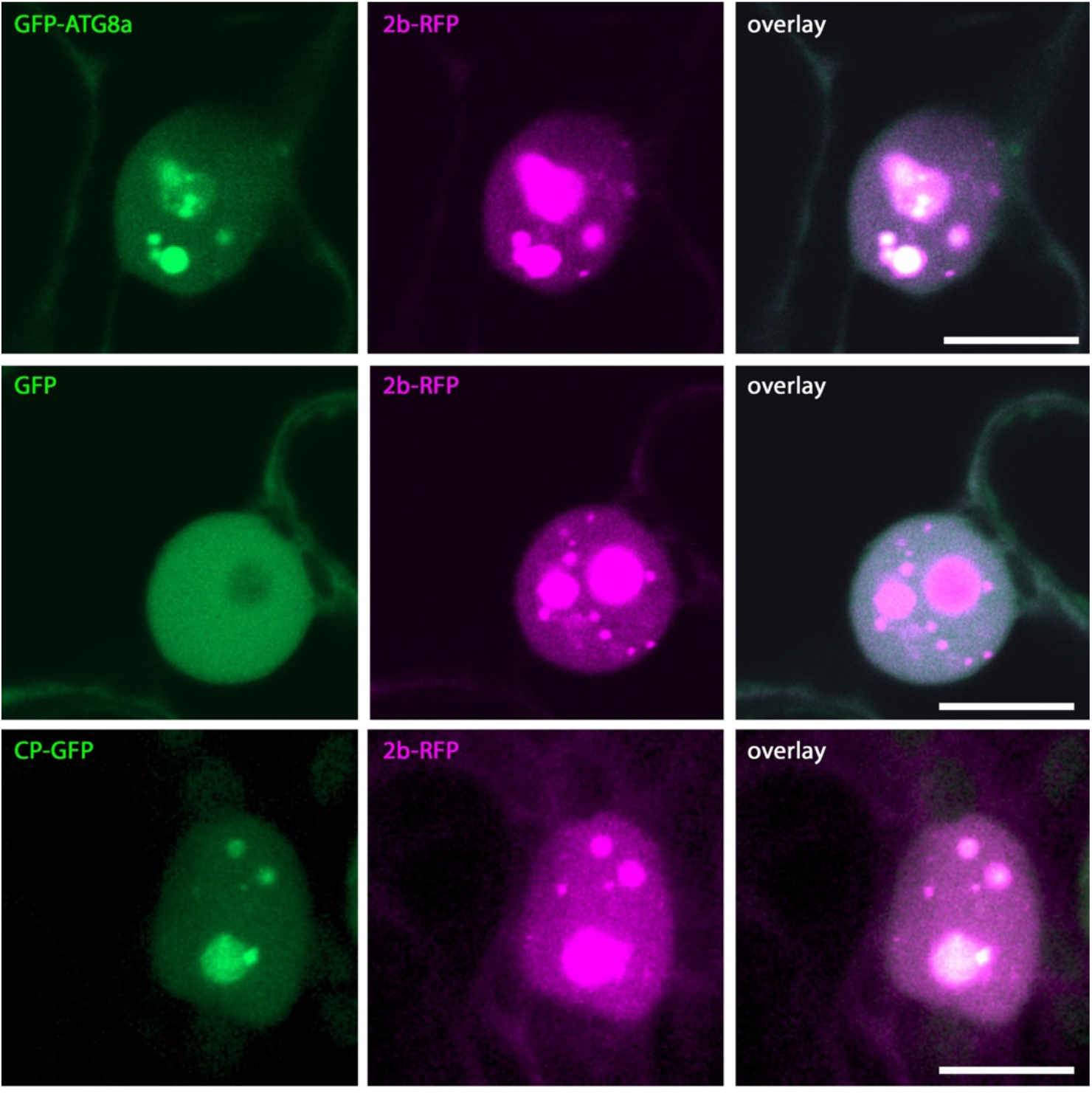
Nuclear localization analysis of 2b-RFP, GFP-ATG8a, GFP and CP-GFP. 2b-RFP was co-expressed with GFP-ATG8a, GFP or CP-GFP in *N. benthamiana* and imaged two days post agroinfiltration. GFP-ATG8a and GFP are single plains and CP-GFP a Z-stack projection. Scale bar = 10 μm.

**S2 Fig.**
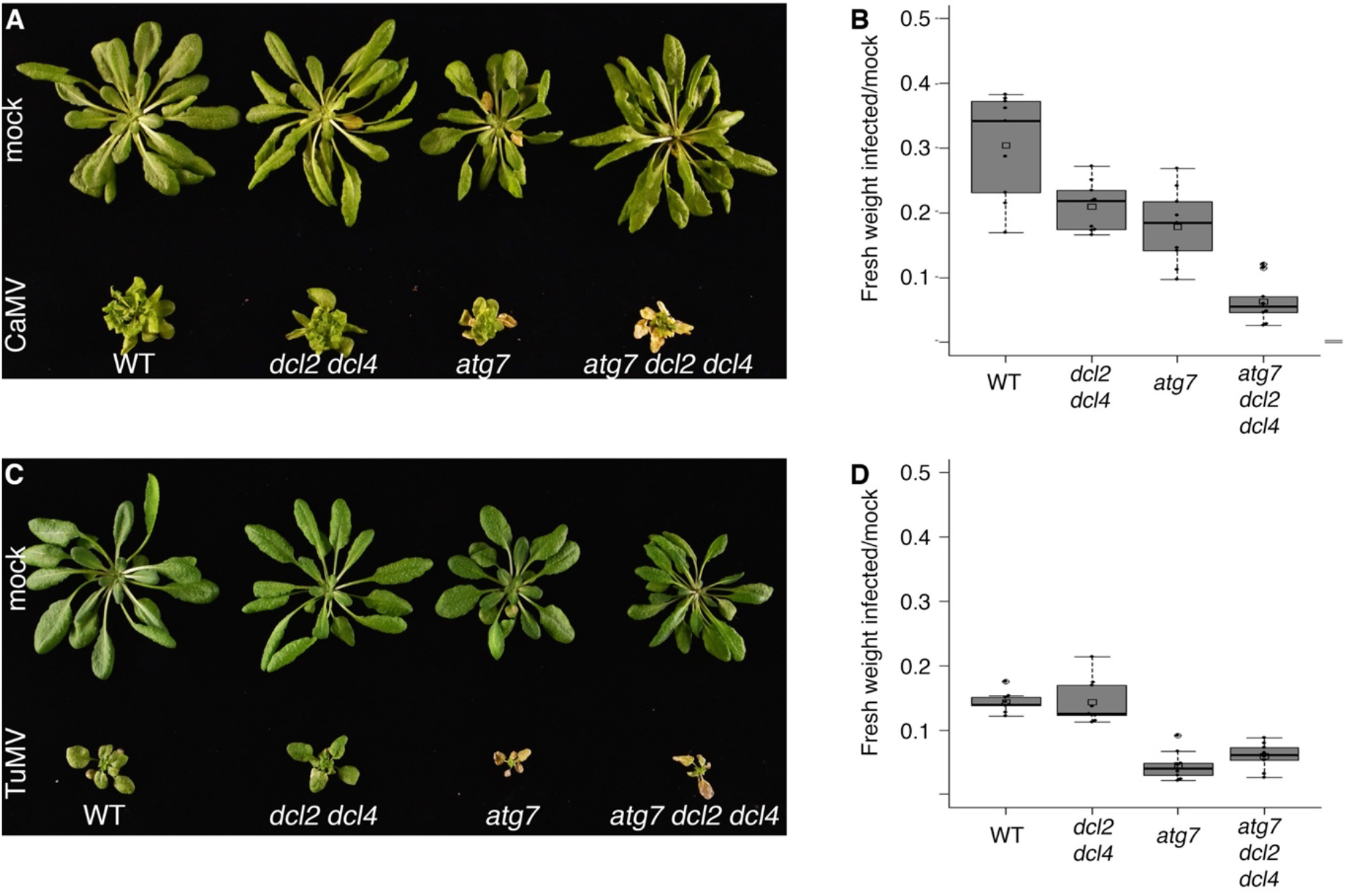
RNA silencing and autophagy provide additive protection against CaMV but not TuMV disease. A) Representative image of mock and CaMV WT, *dcl2 dcl4*, *atg7* and *atg7 dcl2 dcl4* plants at 28 DAI. B) Fresh weight index representing the ratio between infected to mock plants. (*n*=9). C) Representative image of mock and TuMV WT, *dcl2 dcl4*, *atg7* and *atg7 dcl2 dcl4* plants at 21 DAI. D) Fresh weight index representing the ratio between infected to mock plants. (*n*=9).

